# Horn size is linked to Sertoli cell efficiency and sperm size homogeneity during sexual development in common eland (*Taurotragus oryx*)

**DOI:** 10.1101/2024.06.04.597317

**Authors:** Eliana Pintus, Radim Kotrba, José Luis Ros-Santaella

**Author notes:** Correspondence **(EP),** **(JLR-S)**.

## Abstract

**Background:** In polygynous species, the development of secondary sexual characters is usually decisive for male reproductive success. However, our understanding about the links between the growth of these traits and reproductive efficiency is still elusive. Most research efforts in this topic have been also focused on adult males, although the development of some secondary sexual characters, like bovid horns, typically starts after birth, continues during the puberty and in some species, such as the common eland, slows or even stops during adulthood. In this study, we investigated the relationships between horn size and testicular function during sexual development in common elands using a comprehensive approach that considers both spermatogenic and sperm parameters.

**Methods:** Twenty-two non-sexually mature common elands were analyzed for the present study. Horn size was estimated by performing a principal component analysis of horn length, spiral length, and basal circumference. In addition, body mass, testes mass, and gonadosomatic index were assessed. Spermatogenic activity was assessed by cytological and histological analyses. Sperm concentration, morphology, morphometry, and intramale variation in sperm size were evaluated on epididymal sperm samples.

**Results:** We found that bigger horns are associated with increased Sertoli cell efficiency and reduced intramale variation in sperm size. Both parameters were not related to one another while they have shown to be associated with enhanced sperm quality in ungulates. Moreover, horn size was positively linked to the testis mass, sperm concentration, and testicular investment in the seminiferous epithelium. Surprisingly, horn size did not correlate to the percentage of the spermatozoa over the germ cell population nor the percentage of those with normal morphology, which typically increase throughout the male sexual development. It is also remarkable to note that the spiral length and basal circumference were the horn traits most strongly correlated with spermatogenic and sperm parameters as well as those responsible for the sexual dimorphism in this species.

**Conclusions:** Taken together, our results indicate that horn size can be regarded as a good index of male reproductive potential during sexual development and provide insights into the role of secondary sexual characters in sexual selection dynamics.

## Introduction

In sexually dimorphic and polygynous species, male secondary sexual traits are regarded as the hallmark of sexual selection as their expression may influence mating success rendering males more successful in male-male contests (e.g., weapons like tusks, antlers, and horns) and/or more attractive to females (e.g., ornaments like plumage/fur color and manes) (Simmons *et al*., 2017; Snook *et al*., 2013). The question on whether the investment in secondary sexual traits growth might predict male fertility has been a matter of a vibrant debate over the last decades. Several studies have explored the relationship between male secondary sexual characters and sperm traits in a variety of taxa (e.g., crustaceans: Paschoal and Zara, 2022; insects: Rogers *et al*., 2008; fish: Kekäläinen *et al*., 2014; amphibians: Doyle, 2011; birds: Navara *et al*., 2012; mammals: Malo *et al*., 2005a), often finding mixed or weak empirical evidence (Mautz *et al*., 2013). An aspect to be considered is that no study has so far explored the relationship between spermatogenic function and the morphology of secondary sexual characters, inasmuch testis size and sperm quality are commonly regarded as the main determinant of sperm production and competitiveness, respectively. To fill this gap of knowledge, Ramm and Schärer (2014) advocate a more comprehensive approach that, over testis size, also considers the testicular architecture, spermatogenic cell organization and their hierarchical relationships. Testis mass is indeed a vague measure of male investment in the seminiferous epithelium as it also includes connective tissue, smooth muscle, nerves, blood and lymphatic vessels with the fluids herein. Another aspect that has been attracting the attention of evolutionary studies entails the role of intramale variation in sperm size, which has been mostly explored in a small number of mammalian species (i.e., rodents: Varea-Sánchez *et al*., 2014; Šandera *et al*., 2013). To date, only one study has investigated the implications of intramale variation in sperm size in an ungulate species, the red deer (Ros-Santaella *et al*., 2015). In this study, a reduced coefficient of variation (CV) in sperm size was associated with greater testes mass, sperm velocity and normal morphology, but it is still unknown whether the homogeneity in the sperm size is associated with the development of secondary sexual characters. Additionally, most research efforts exploring the relationships between pre- and post-copulatory traits in mammals have been so far focused on sexually mature males (Dines *et al*. 2015; Ferrandiz-Rovira *et al*. 2014), even though the development of secondary sexual characters in many species (e.g., some bovids) starts early in the male lifetime and sometimes even do not appreciably grow during adulthood (Geist, 1966).

In several bovids and cervids, horns and antlers are one of the most diverse and elaborate male secondary sexual characters. In addition to their role as sexual traits, horns and antlers play a major role as weapons against predators and temperature regulators (Bro-Jørgensen, 2007; Picard *et al*., 1999). Previous studies have shown that horn/antler size and shape in male ungulates are associated with fighting behavior (Caro, 2003; Lundrigan, 1996), major histocompatibility complex (MHC) traits (Ditchkoff *et al*., 2001), parasite abundance (Ezenwa and Jolles, 2008), testis size (Malo *et al*., 2005a), sperm motility and velocity (Santiago-Moreno *et al*., 2007; Malo *et al*., 2005a), and reproductive success (Willisch *et al*., 2015; Preston *et al*., 2003; Kruuk *et al*., 2002). In a comparative study in ungulates, Ferrandiz-Rovira *et al*. (2014) found that longer weapons were significantly associated with shorter sperm cells, but there was no association between weapons length and testes mass (but see also Lüpold *et al*., 2015). In addition to their size, also fluctuating asymmetry in bilateral paired organs, like horns, can be used as an index of developmental stability (Benítez *et al*., 2020). In ungulates, fluctuating asymmetry in male secondary sexual traits has been related to poor ejaculate quality, at least in captive populations (Roldan *et al*., 1998). Surprisingly, the links between horn size/asymmetry and testicular function are still largely unknown in ungulates.

The aim of this study was to explore the relationships between horn morphology (i.e., size and asymmetry) and reproductive competence during the sexual development of male common eland (*Taurotragus oryx*). The common eland is one of the largest antelope species native to the southern and eastern Africa. This non-territorial and sexually dimorphic species is currently classified as “least concern” by the IUCN, with a stable population of ∼100,000 individuals (IUCN, 2024). The age of sexual maturity is estimated to be around 2.5 years in females and 4 years in males (Pappas, 2002). Elands can reproduce at any time of the year, although in wild conditions breeding and calving season peak might occur during the rainy season (Pennigton 2009; Pappas, 2002). Both sexes have spiral horns, which are shorter, thicker, and have tighter and more pronounced spirals in males than in females (Pappas, 2002). Final horn length is achieved relatively early during male development and do not appreciable grow after reaching sexual maturity (Jeffery and Hanks, 1981). For this reason, sexual development can represent a crucial phase of the male lifetime in which a link between primary and secondary characters is established. The understanding of the role played by horn morphology during the male sexual development can reveal early important traits of individual fitness that might be determinant for reproductive success during adulthood.

## Materials and Methods

### Animal management and experimental design

From September 2013 to December 2017, 22 male common elands (age: 15-44 months old; **Figure 1A**) were slaughtered at the farm of the Czech University of Life Sciences Prague (Lány, Czech Republic). The farm is accredited as research facility according to European and Czech laws for ethical use of animals in research (permits no. 58176/2013-MZE-17214 and no.

**Figure 1.**
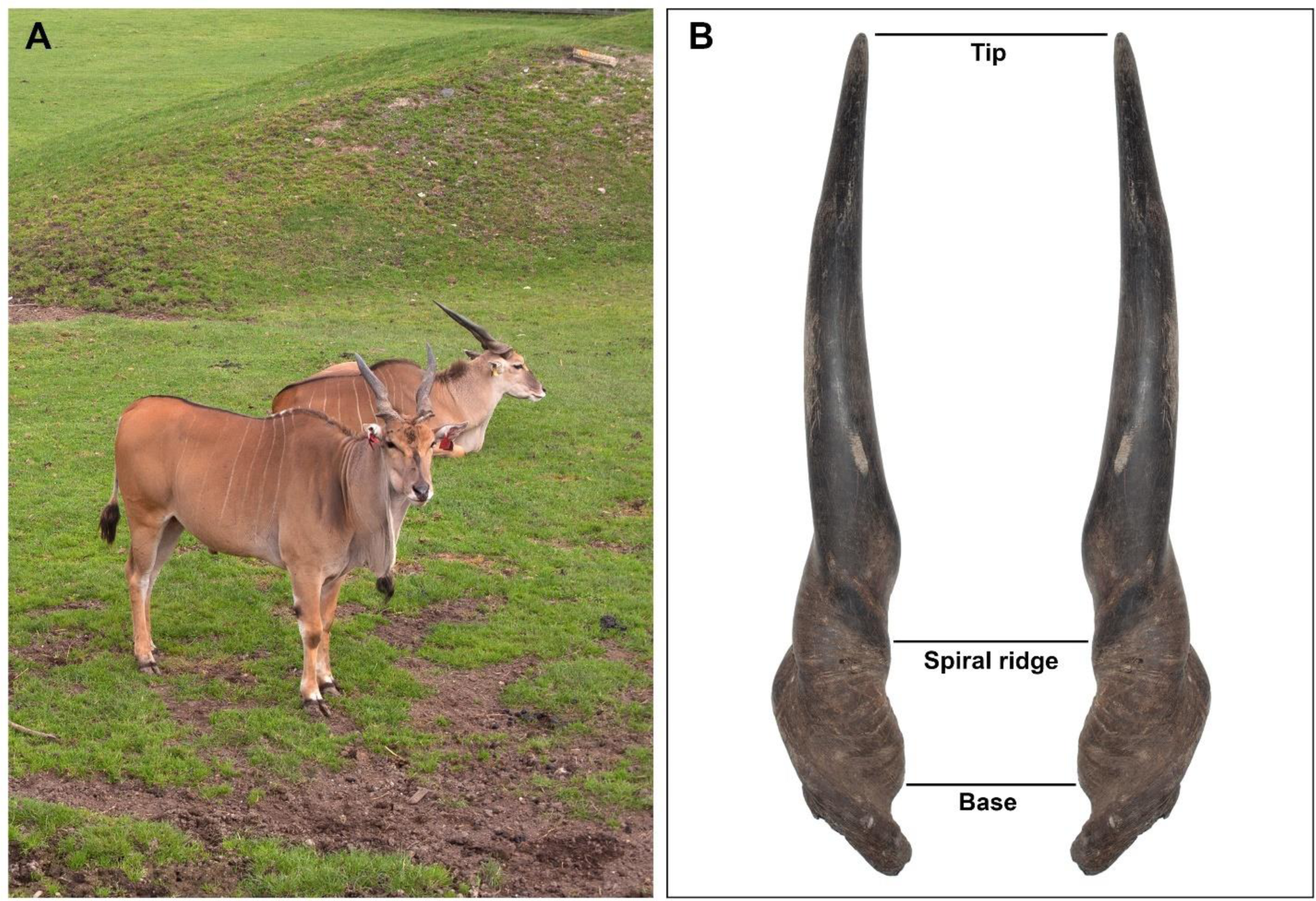
Post-pubertal males and horn measurements in common eland. A) Post-pubertal male common elands. B) Specular image that representatively shows the points of reference for horn size measurements: horn length (from the front base and straight up to the tip), spiral length (from the front base following the spiral ridge to the point where the latter is not pronounced and then straight up to the tip), and basal circumference.

63479/2016-MZE-17214). Slaughter of males is part of farm regular production and management to reduce the number of animals due to overwintering capacity. All individuals were born in captivity, individually identified by ear tags since birth, and bred under the same environmental conditions. They represented the seventh captive generation after their import from East Africa from 1969 to 1972 (Vágner, 1974). Genetic diversity of the herd has been maintained through exchanges of individuals from zoological gardens. All eland males were group-housed in straw bedded pens in a barn together with the rest of the herd of maximum of 50 animals, which was separated usually into two groups based on reproductive state of adult females and time of year. All animals were fed with mixed diets *ad libitum* consisted of corn silage (60%), lucerne haylage (30%), meadow hay (7%), and barley straw (3%). This mixture contained 16.6% of crude protein and 16.2% of crude fiber. From April to November, they had access to 2.5 ha of paddock with grass to enhance contact between animals and received up to one kg/individual/day of barley grain. The health of animals was randomly checked by veterinarian inspection and monitored on yearly basis. The animals were slaughtered, exsanguinated, eviscerated at the farm, and transported to the abattoir of the Institute of Animal Science in Prague for further processing (see more in Bartoň *et al*., 2014). All slaughter process was carried out under the supervision of a state veterinarian according to EU and national legislation and conditions for farm animals (slaughter permit no. SVS/WS22/2012-KVSS and no. SVS/2015/077267-S). Testes (within the scrotum) were removed directly after the slaughter, stored in sealed plastic bags, labelled according to individual animal recognition, and transported at room temperature to the laboratory for further processing. Testicular and sperm samples were collected and processed within approximately 2 hours after the death of the animals.

### Biometrics

Body mass was determined using tensometric scale (EC2000, True-Test Limited, Auckland, New Zealand) to the nearest 0.5 kg. Horn size was determined by flexible tape measure to the nearest 0.5 cm. Horn size measurements were determined as it follows: horn length (from the front base and straight up to the tip), spiral length (from the front base following the spiral ridge to the point where it is not pronounced and then straight up to the tip), and basal circumference (**Figure 1B**). All horn measurements were taken by the same trained observer (RK) and averaged from the left and right sides. Horn asymmetry was calculated as the signed difference between right and left sides of each trait (Palmer, 1994). Horn size was not adjusted for body size as scaling relationship between organs typically occurs during ontogenetic growth (Vea and Shingleton, 2021). In comparative biology, recent studies have also questioned traditional methods for body-size adjustment as they i) do not adequately separate the effects of body size from those of other biological and ecological factors on a specific phenotypic trait (Glazier, 2022) and ii) can spuriously change the sign of regression coefficients compared to the original values, which could lead to inferential biases in biological studies (Rogell *et al*., 2019).

### Testes mass and spermatogenic function

Testes mass was recorded to the nearest 0.1 g using an electronic balance (EK-600G, LTD, Japan). The GSI was calculated as the relative proportion of testes mass to the body mass. Cytological samples were collected from each testis using the fine needle aspiration technique, which has proved to be a reliable method for the assessment of testicular function in ungulates both under physiological and pathological conditions (Pintus *et al*., 2014, 2015a). Testicular smears were stained with Hemacolor (Merck, Darmstadt, Germany) and evaluated under a 100× objective using bright-field microscopy (Nikon Eclipse E600, Nikon, Tokyo, Japan). Assessment of germ cell proportions and spermatogenic indices were determined on at least 200 Sertoli and spermatogenic cells per testis (**Figure 2A**). Then, testicular indices were assessed as follows: i) the Sertoli cell index (SEI), which is the percentage of Sertoli cells per total germ cells and estimates the spermatogenic activity; ii) the Spermatozoa-Sertoli cell index (SSEI), which is the number of spermatozoa per Sertoli cell; iii) the meiotic index (MI), which is the ratio of round spermatids to primary spermatocytes, and estimates the germ cell loss during meiosis; iv) the ratio of elongated spermatids to round spermatids (ES/RS), which estimates the germ cell loss during the post-meiotic phase; v) the ratio of elongated spermatids to total germ cells (ES/GC), which estimates the overall germ cell loss during spermatogenesis; vi) the ratio of round spermatids to Sertoli cells (RS/SC), vii) the ratio of elongated spermatids to Sertoli cells (ES/SC), which both estimate the Sertoli cell function; and viii) the ratio of total germ cells per Sertoli cells (GC/SC) that estimates the Sertoli cell workload capacity (Ros-Santaella *et al*., 2019). Samples for testicular histology were collected and processed as previously described (Pintus *et al*., 2015a). To evaluate the morphology of seminiferous tubules, 25 roundish cross-sections of the seminiferous tubules were photographed per each testis using a high-resolution camera (Digital Sight DSFi1, Nikon, Tokyo, Japan) under a 20× objective. Then, the area of the seminiferous tubule, lumen, and epithelium were assessed using ImageJ software (National Institutes of Health, Bethesda, MD, USA) (**Figure 2B**). The proportion of the seminiferous epithelium to the total tubular area was then calculated. Values from cytological and histological analyses were averaged from the left and right testes. Because of suboptimal sample quality or reduced smear cellularity, testicular cytology and histology were not performed in one and two males, respectively. All measurements were taken by the same trained observer (EP).

**Figure 2.**
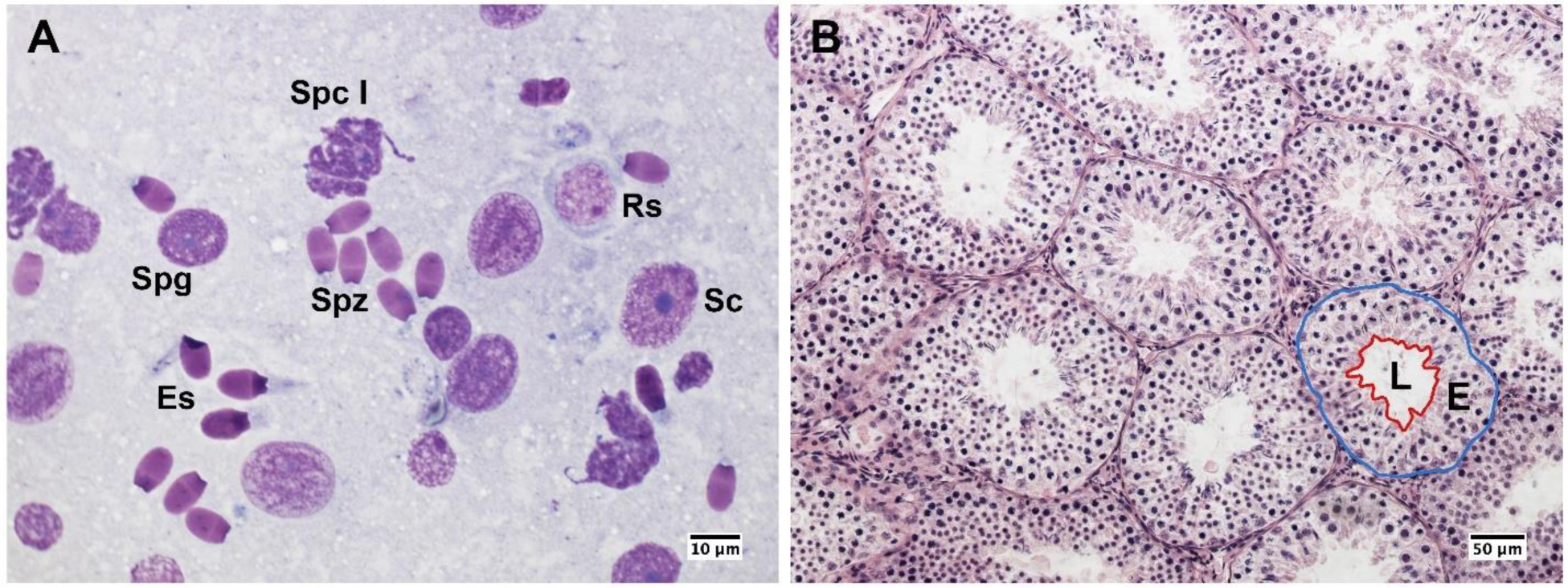
Testicular cytology and histology from post-pubertal common eland. A) Germ cells and Sertoli cells from a cytological smear. Sc: Sertoli cell; Spg: spermatogonium; Spc I: Primary spermatocyte; Rs: round spermatid; Es: elongated spermatid; Spz: spermatozoon. B) Histological section of seminiferous tubules that shows a representative example of seminiferous tubule measurements: the blue line indicates the area of the seminiferous tubule, whereas the red line indicates the epithelial (E) and luminal (L) areas.

### Sperm concentration, morphology, and morphometry

Sperm samples were collected from the epididymal caudae using a sterile surgical blade and fixed into 0.5 mL phosphate-buffered saline solution supplemented with 2% glutaraldehyde. Sperm concentration was determined using a Bürker chamber. The percentage of morphologically normal spermatozoa was determined after evaluating 200 spermatozoa under phase-contrast microscopy (40× objective). Sperm morphometry was assessed as previously described (Ros-Santaella *et al*., 2014, 2015). Briefly, sperm pictures were taken with high-resolution camera under phase-contrast microscopy (40× objective). The following sperm traits were determined using the ImageJ software: head width, head length, midpiece length, and flagellum length. From these measurements, other morphometric parameters were calculated such as head area, head perimeter, head ellipticity (head length/head width), total sperm length, and principal plus terminal piece length. Twenty-five spermatozoa with normal morphology (i.e., without head or flagellum abnormalities and without proximal cytoplasmic droplet) were measured per male. For each parameter, the intramale coefficient of variation (CV) was calculated as standard deviation/mean×100. The main structures of common eland spermatozoa with normal morphology are shown in **Figure 3**. Sperm morphology and morphometry could not be assessed in five samples because of their low sperm concentration on the smears. All measurements were taken by the same trained observer (JLR-S).

**Figure 3.**
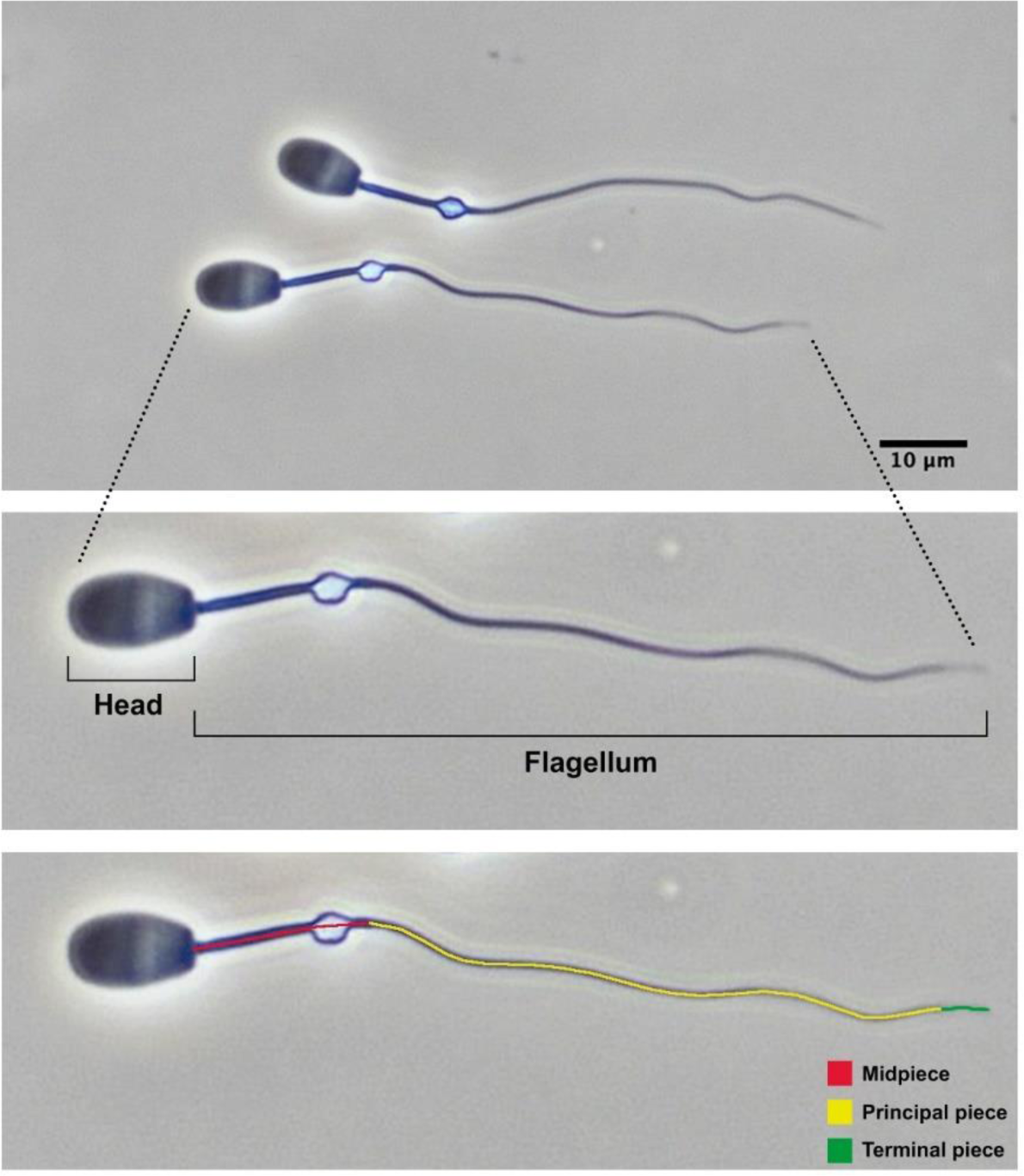
Epididymal spermatozoa from post-pubertal common eland and their main structures.

### Statistical analysis

Statistical analyses were performed using the SPSS 20.0 statistical software (IBM Inc., Chicago, IL, USA). The Shapiro-Wilk test was used to check for the normal distribution of data. Data that were not normally distributed were log-transformed. Non-parametric tests were applied to data that were not normally distributed after log transformation. Principal component analysis was used to reduce several closely related variables (i.e., horn size/asymmetry, Sertoli cell efficiency, and intramale variation in sperm head size) into a smaller subset that better summarizes the original data. The Bartlett sphericity and Keiser-Meyer-Olkin (KMO) tests for sampling adequacy were applied to test the suitability of data for the principal component analysis. The principal components (PCs) with Bartlett sphericity test’s *p* value higher than 0.05 and KMO test lower than 0.5 were considered inappropriate (Budaev, 2010). Two-tailed Pearson correlations were used when data were normally distributed, otherwise two-tailed Spearman correlations.

Correlation analyses were not corrected for multiplicity because, while correction methods decrease the probability of Type I error, they increase the probability of Type II error (Streiner, 2015). Therefore, although the statistical significance was set at *p*<0.05, *p* values close to 0.05 must be cautiously considered. Data are shown as the mean±SE.

## Results

Descriptive statistics of biometrics, testicular, and sperm parameters of male common elands are shown in **Tables 1** and **2**. On average, horn size was around 55 cm in length and 26 cm at basal circumference, while the spiral length was 68 cm. The signed asymmetry between the right and left sides of each horn trait was: −0.68±0.36 cm for horn length, −0.64±0.46 cm for spiral length, and −0.14±0.19 cm for basal circumference. Except one male in which testicular cytology could not be assessed although spermatozoa were present in the epididymal caudae, the spermatogenesis was complete in all individuals, which indicate that males did already reach puberty. Cytological and histological analyses showed that spermatozoa represent around one fifth of spermatogenic cell population, with over 80% of the tubular area filled by the seminiferous epithelium. Among spermatogenic cells, the round spermatids were the most abundant, while secondary spermatocytes were the scarcest. On average, over 700×10^6^ spermatozoa/mL were collected from the epididymal caudae, with two thirds of them being morphologically normal. The proportion of each sperm structure in relation to the total sperm length was: head length, 13.45%; midpiece length, 20.21%; and principal piece plus terminal piece length, 66.34%. Overall, size variation in sperm parameters was below 5%, relatively higher CV values were found in the sperm head than in the flagellum (**Table 2**).

**Table 1.** Descriptive statistics of biometrics and testicular parameters in post-pubertal common eland.

|  | Mean±SE | N |
| --- | --- | --- |
| *Biometrics* |  |  |
| Body mass (kg) | 273.25±16.23 | 22 |
| Horn length (cm) | 55.23±1.07 | 22 |
| Horn spiral length (cm) | 67.82±1.50 | 22 |
| Horn basal circumference (cm) | 25.89±0.29 | 22 |
| Testes mass (g) | 123.43±7.28 | 22 |
| GSI (%) | 0.05±0.00 | 22 |
| *Spermatogenic cell subtypes* |  |  |
| Spermatogonia (%) | 2.25±0.25 | 21 |
| Primary spermatocytes (%) | 22.65±0.83 | 21 |
| Secondary spermatocytes (%) | 0.88±0.10 | 21 |
| Round spermatids (%) | 37.51±0.69 | 21 |
| Elongated spermatids (%) | 16.76±0.67 | 21 |
| Spermatozoa (SI, %) | 19.95±0.76 | 21 |
| *Spermatogenic indices* |  |  |
| SEI (%) | 11.44±1.96 | 21 |
| SSEI | 2.51±0.33 | 21 |
| MI | 1.72±0.08 | 21 |
| ES/RS | 0.46±0.02 | 21 |
| ES/GC | 0.17±0.01 | 21 |
| RS/SC | 4.74±0.68 | 21 |
| ES/SC | 2.05±0.25 | 21 |
| GC/SC | 12.37±1.61 | 21 |
| *Testicular architecture* |  |  |
| Area seminiferous tubule (µm<sup>2</sup>) | 26,520.26±1,050.76 | 20 |
| Area seminiferous epithelium (µm<sup>2</sup>) | 22,033.46±936.32 | 20 |
| Area seminiferous lumen (µm<sup>2</sup>) | 4,486.80±219.69 | 20 |
| Proportion of the seminiferous epithelium to the tubular area (%) | 83.07±0.82 | 20 |

GSI: Gonadosomatic index; SI: spermatogenic index; SEI: Sertoli cell index; SSEI: Spermatozoa-Sertoli cell index; MI: meiotic index; ES/RS: elongated spermatids per round spermatids ratio; ES/GC: elongated spermatids per germ cells ratio; RS/SC: round spermatids per Sertoli cells ratio; ES/SC: elongated spermatids per Sertoli cells ratio; GC/SC: germ cells per Sertoli cells ratio. Data are shown as mean±SE.

**Table 2.** Descriptive statistics of sperm parameters in post-pubertal common eland.

|  | Mean±SE | N |
| --- | --- | --- |
| *Sperm number and morphology* |  |  |
| Sperm concentration (10<sup>6</sup>/mL) | 755.32±100.76 | 21 |
| Normal sperm (%) | 68.79±7.01 | 17 |
| *Sperm size* |  |  |
| Head width (µm) | 5.92±0.04 | 17 |
| Head length (µm) | 9.66±0.07 | 17 |
| Head perimeter (µm) | 24.84±0.13 | 17 |
| Head area (µm<sup>2</sup>) | 44.95±0.44 | 17 |
| Head ellipticity | 1.63±0.02 | 17 |
| Midpiece length (µm) | 14.52±0.10 | 17 |
| Principal plus terminal piece length (µm) | 47.66±0.38 | 17 |
| Flagellum length (µm) | 62.17±0.42 | 17 |
| Sperm length (µm) | 71.84±0.43 | 17 |
| *Intramale CV in sperm size* |  |  |
| Head width (%) | 3.29±0.20 | 17 |
| Head length (%) | 2.85±0.15 | 17 |
| Head perimeter (%) | 2.21±0.10 | 17 |
| Head area (%) | 4.35±0.21 | 17 |
| Head ellipticity (%) | 4.46±0.20 | 17 |
| Midpiece length (%) | 2.68±0.16 | 17 |
| Principal plus terminal piece length (%) | 2.07±0.08 | 17 |
| Flagellum length (%) | 1.57±0.07 | 17 |
| Sperm length (%) | 1.49±0.06 | 17 |

CV: coefficient of variation. Data are shown as mean±SE.

### Principal component analysis of horn size and asymmetry

A principal component analysis was performed using the averaged values of the left and right horn measurements to obtain smaller subset of variables that summarizes the horn size. We obtained a single component (PC1, horn size) that overall explained 75.89% of the total variance (Keiser-Meyer-Olkin, KMO, measure of sampling adequacy=0.520; Bartlett’s test of sphericity: approx. χ^2^=44.41, df=3, *p*<0.0001; **Table 3**) and showed normal distribution (Shapiro-Wilk test, *p*=0.872). Another principal component analysis was performed using the signed difference between the right and left horn measurement to obtain a smaller subset of variables that explain horn asymmetry. We obtained a single component (PC1, horn asymmetry) that overall explained 60.29% of the total variance (KMO measure of sampling adequacy=0.516; Bartlett’s test of sphericity: approx. χ^2^=16.30, df=3, *p*=0.001; **Table 3**). The PC1 of horn asymmetry shows normal distribution (Shapiro-Wilk test, *p*=0.552) around a mean value close to zero (Skewness=-0.554±0.491 and Kurtosis=0.496±0.953), which is indicative of fluctuating asymmetry.

**Table 3.** Principal component analysis of horn size and asymmetry in post-pubertal common eland.

|  | PC1 | *p* |
| --- | --- | --- |
| *Horn size* |  |  |
| Horn length | 0.916 | <0.001 |
| Spiral length | 0.967 | <0.001 |
| Basal circumference | 0.709 | <0.001 |
| Eigenvalue | 2.28 |  |
| Variance explained (%) | 75.89 |  |
| *Horn asymmetry* |  |  |
| Horn length | 0.923 | <0.01 |
| Spiral length | 0.909 | <0.01 |
| Basal circumference | 0.361 | <0.05 |
| Eigenvalue | 1.81 |  |
| Variance explained (%) | 60.29 |  |

PC: Principal component. Horn asymmetry was calculated as the signed value of the difference between right and left sides of each trait.

### Correlations between horn size and reproductive function

#### Correlations of horn size with testicular and spermatogenic parameters

We found that horn size was significantly associated with higher testes mass (r=0.476 *p*=0.025), while it did not correlate with the gonadosomatic index (GSI log-transformed; r=0.083 *p*=0.713). Albeit no significant, horn size was positively associated with the body mass (log-transformed; r=0.421 *p*=0.051). An unexpected finding was that horn size was not related to the proportion of any spermatogenic cell type, including the spermatic index that represents the proportion of spermatozoa over the total germ cells and estimates the sperm production (r=0.071, *p*=0.761). On the other hand, larger horn size was associated with reduced Sertoli cell index (SEI, log-transformed r=-0.590, *p*=0.005) and increased Sertoli cell function and workload capacity (i.e., Spermatozoa-Sertoli cell index, SSEI: r=0.531, *p*=0.013; ratio of round spermatids per Sertoli cell, RS/SC: r=0.540, *p*=0.012; ratio of elongated spermatids per Sertoli cell, ES/SC: ρ=0.516, *p*=0.017; ratio of total germ cells per Sertoli cell, GC/SC: r=0.581, *p*=0.006, all log-transformed). Among the three descriptors of horn size, spiral length was the one showing the strongest relationship with Sertoli cell function and workload (i.e., Spermatozoa-Sertoli cell index, SSEI: r=0.603, *p*=0.004; ratio of round spermatids per Sertoli cell, RS/SC: r=0.610, *p*=0.003; ratio of elongated spermatids per Sertoli cell, ES/SC: ρ=0.527, *p*=0.014; ratio of total germ cells per Sertoli cell, GC/SC: r=0.650, *p*=0.001, all log-transformed). To corroborate our findings, we performed a principal component analysis of Sertoli cell indices because of the high correlation among them. We obtained a single principal component (PC1, Sertoli cell efficiency) that explained 96.74% of the total variance (KMO measure of sampling adequacy=0.803; Bartlett’s test of sphericity: approx. χ^2^=268.75, df=10, *p*<0.0001; **Table 4**) and showed normal distribution (Shapiro-Wilk test, *p*=0.055). Our findings confirm that larger horn size was associated with increased Sertoli cell efficiency (r=0.589, *p*=0.005, **Figure 4**).

**Figure 4.**
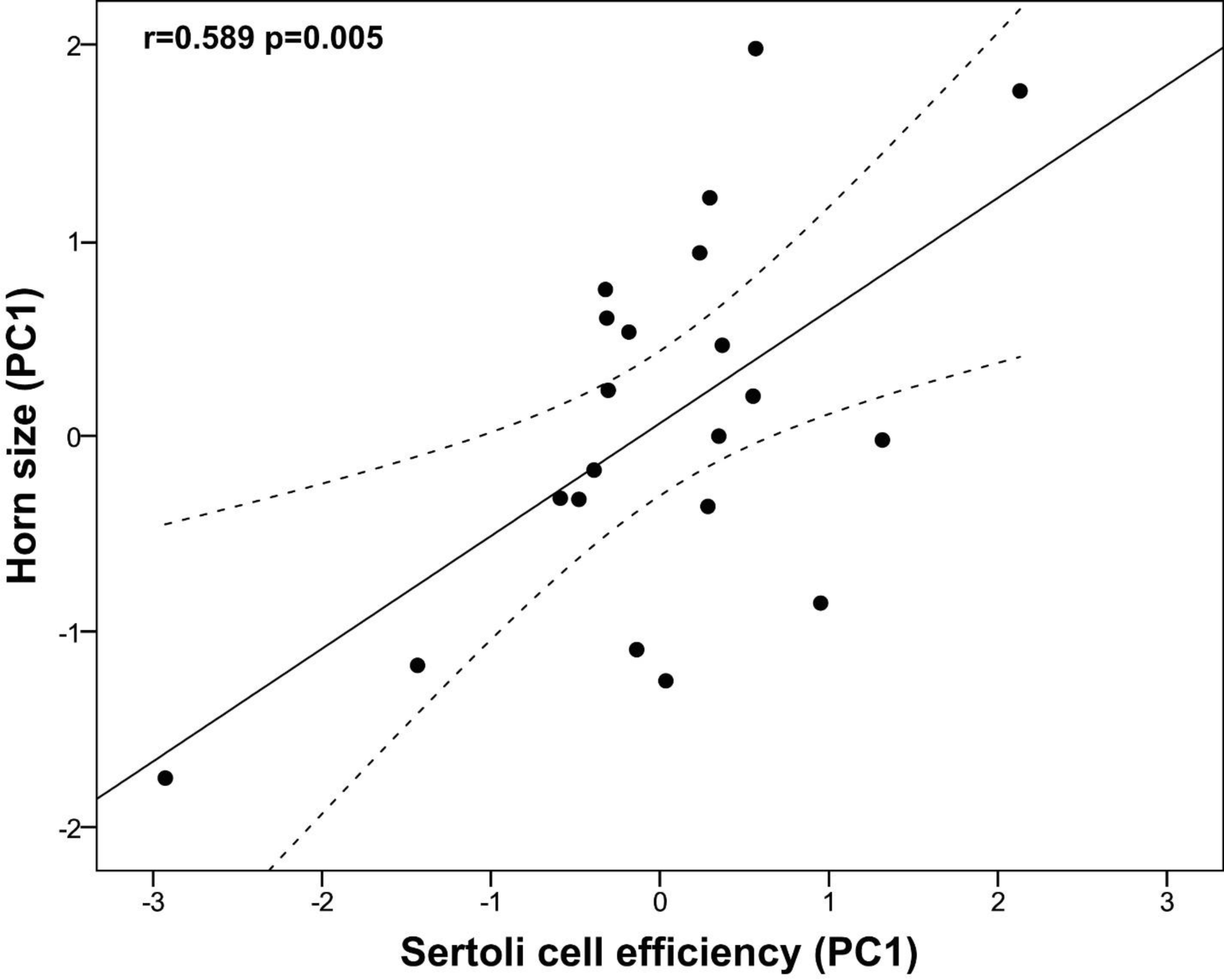
Relationship between horn size and Sertoli cell efficiency in post-pubertal common eland. Black line represents linear fit line, while dashed lines indicate 95% confidence intervals. PC: Principal component.

**Table 4.** Principal component analysis of Sertoli cell efficiency and intramale variation in sperm head size in post-pubertal common eland.

|  | PC1 | *p* |
| --- | --- | --- |
| *Sertoli cell efficiency* |  |  |
| Log SEI | -0.997 | <0.001 |
| Log SSEI | 0.968 | <0.001 |
| Log RS/SC | 0.986 | <0.001 |
| Log ES/SC | 0.968 | <0.001 |
| Log GC/SC | 0.998 | <0.001 |
| Eigenvalue | 4.84 |  |
| Variance explained (%) | 96.74 |  |
| *Intramale variation in sperm head size* |  |  |
| Log head width CV | 0.938 | <0.001 |
| Head area CV | 0.937 | <0.001 |
| Head ellipticity CV | 0.901 | <0.001 |
| Eigenvalue | 2.57 |  |
| Variance explained (%) | 85.64 |  |

CV: coefficient of variation; SEI: Sertoli cell index; SSEI: Spermatozoa-Sertoli cell index; RS/SC: rounds spermatids per Sertoli cells ratio; ES/SC: elongated spermatids per Sertoli cells ratio; GC/SC: germ cells per Sertoli cells ratio. PC: Principal component.

Horn size was also positively related to the area of the seminiferous tubule and epithelium (r=0.462, *p*=0.040 and r=0.531, *p*=0.016, respectively). There was also a significant positive relationship between horn size and the proportion of the tubular area lined by the seminiferous epithelium (ρ=0.456, *p*=0.043). Among the three descriptors of horn size, basal circumference was the one showing the strongest relationship with the area of seminiferous tubules and epithelium (r=0.640, *p*=0.002 and r=0.629, *p*=0.003, respectively). We performed a principal component analysis of testicular architecture, which rendered one principal component that explained 77.13% of the total variance. However, because of the low value of KMO measure of sampling adequacy test (i.e., 0.327), the PC was rejected.

#### Correlations of horn size with sperm parameters

Horn size was positively associated with sperm concentration (r=0.543, *p*=0.011) but it did not correlate with either sperm morphology or size (*p*>0.05). Albeit not significant, large horn size was associated with a reduced midpiece length (r=-0.481, *p*=0.051). Interestingly, bigger horn size was significantly associated with reduced intramale variation in sperm head size (log-head width CV: r=-0.614, *p*=0.009; head area CV: r=-0.520, *p*=0.033) and shape (head ellipticity CV: r=-0.602, *p*=0.011). Among the three descriptors of horn size, basal circumference was the one showing the strongest relationship with intramale CV of sperm head size and shape (i.e., log-head width CV: r=-0.549, *p*=0.023; head area CV: r=-0.645, *p*=0.005; head ellipticity CV: r=-0.664, *p*=0.004). To corroborate our findings, a principal component analysis of sperm head CV parameters was performed. We obtained a single principal component (PC1, intramale variation in sperm head size) that explained 85.64% of the total variance (KMO measure of sampling adequacy=0.746; Bartlett’s test of sphericity: approx. χ^2^=31.275, df=3, *p*<0.0001; **Table 4**) and showed normal distribution (Shapiro-Wilk test, *p*=0.234). Our findings support that larger horn size is associated with reduced intramale variation in sperm head size and shape (r=-0.625, *p*=0.007, **Figure 5**).

**Figure 5.**
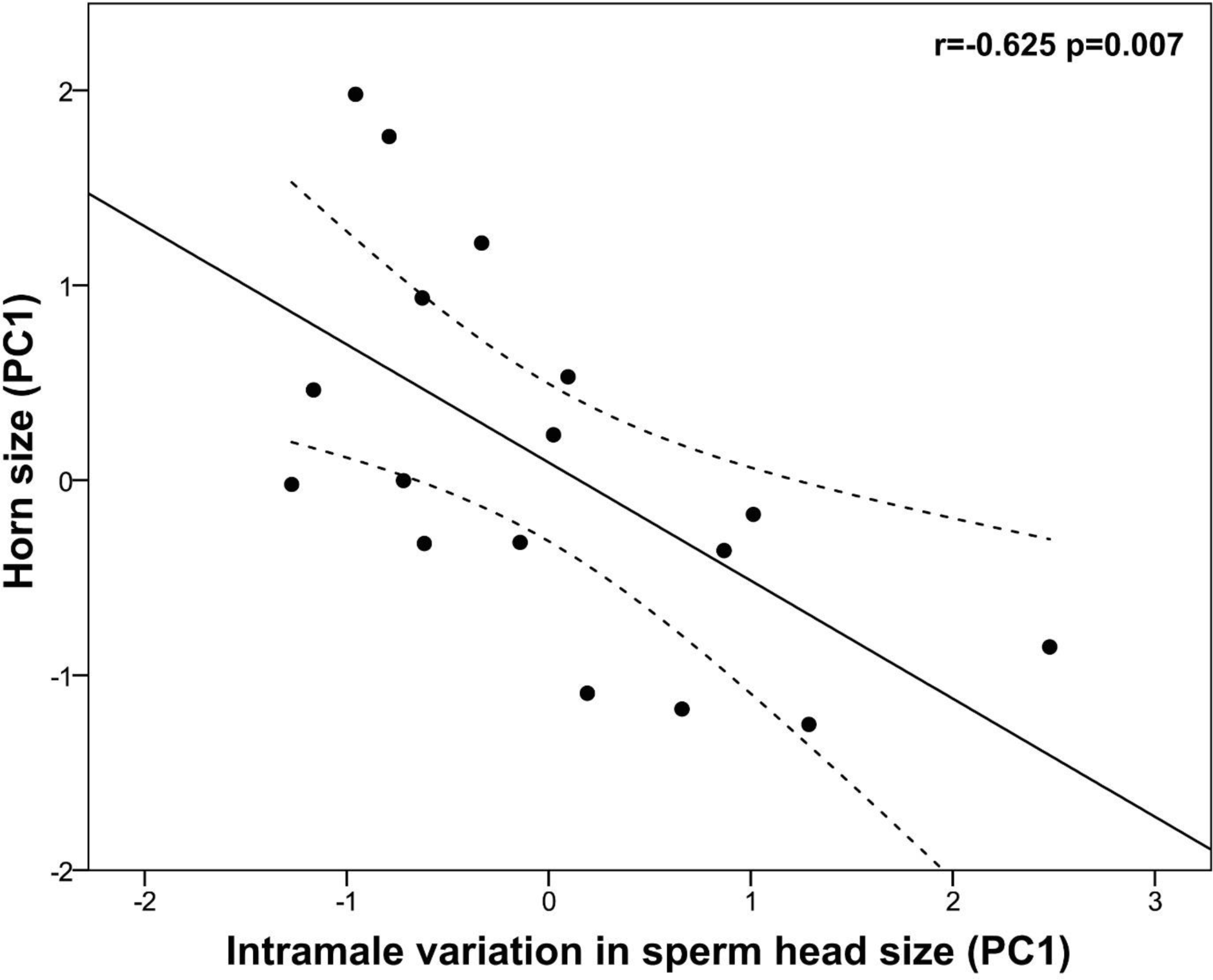
Relationship between horn size and sperm head size homogeneity in post-pubertal common eland. Black line represents linear fit line, while dashed lines indicate 95% confidence intervals. PC: Principal component.

### Correlations between spermatogenic and sperm parameters

To exclude collinearity, we checked for correlation between spermatogenic and sperm parameters that have previously shown to be associated with horn size. We found that increased Sertoli cell efficiency was associated with greater testes size (r=0.652, *p*=0.001), sperm concentration (r=0.475, *p*=0.034), and investment in tubular and epithelial areas of the testicular parenchyma (r=0.515, *p*=0.024 and r=0.587, *p*=0.008, respectively). Surprisingly, reduced intramale variation in sperm head size was not significantly associated either with testes mass (r=-0.167, *p*=0.522), sperm concentration (r=-0.391, *p*=0.121) or Sertoli cell efficiency (r=-0.248, *p*=0.338). However, low intramale variation in sperm head ellipticity was associated with greater sperm concentration (r=-0.500, *p*=0.041) and larger areas of the seminiferous tubule and epithelium (r=-0.590, *p*=0.021 and r=-0.591, *p*=0.020, respectively).

We also checked the correlations between the remaining spermatogenic and sperm parameters (for simplicity, only those with *p*<0.01 are shown herein). We found that high Sertoli cell efficiency was associated with smaller sperm head (i.e., head width: r=-0.659, *p*=0.004; head area: r=-0.678, *p*=0.003), while high meiotic index was strongly associated with shorter sperm length (i.e., principal plus terminal piece length: r=0.834, *p*<0.0001; flagellum length: r=0.822, *p*<0.0001; total sperm length: r=0.819, *p*<0.0001). Sperm concentration was positively associated with tubular and epithelial areas of the testicular parenchyma (r=0.756, *p*<0.001 and r=0.771, *p*<0.001, respectively). Moreover, greater proportion of normal spermatozoa was associated with high Sertoli cell efficiency (ρ=0.725, *p*<0.001) and reduced intramale CV in sperm length (flagellum length: ρ=-0.635, *p*=0.006; total sperm length: ρ=-0.673, *p*=0.003).

### Correlations between horn asymmetry and reproductive function

We found that horn asymmetry (PC1, horn asymmetry) was not significantly correlated neither with biometric, testicular or sperm parameters (*p*>0.05).

## Discussion

The present study comprehensively explores the relationships between primary and secondary sexual characters during the sexual development of a polygynous ungulate, the common eland. We provide evidence that large horn size is associated with increased testis mass, spermatogenic activity, Sertoli cell efficiency, sperm concentration and reduced intramale variation in sperm size. It is remarkable to point out that the strength of correlations between horn size and Sertoli cell efficiency and between the former and sperm size homogeneity showed r>0.58 and *p*<0.01, which provide new insights into the links between primary and secondary sexual characters in a sexually dimorphic mammalian species. Taken together, our results indicate that, during sexual development, horn size is a good index of reproductive potential in the common eland. It remains however to explore whether increased Sertoli cell efficiency and sperm size homogeneity translate into enhanced male fertility.

Our findings confirm the positive relationship between secondary sexual traits and testis size in ungulates (Malo *et al*., 2005a) and suggest that such a relationship is mainly supported by the increased Sertoli cell efficiency. Discovered by Enrico Sertoli in 1865, the Sertoli cell represents the only somatic cell found within the seminiferous tubules. The astonishing ability of Sertoli cells to nurse, protect, and support the development of up to five different types of germ cells at any one time make them one of the most complex cells in the body (O’Donnell *et al*., 2022). The role of Sertoli cells is crucial during the entire male lifetime: during the embryo and foetal development Sertoli cells promote the sexual differentiation, while during puberty and sexual maturity they coordinate the spermatogenesis and determine the sperm production. Our findings also indicate that greater Sertoli cell efficiency is associated with higher percentage of normal spermatozoa and smaller sperm head size, possibly because of Sertoli cell’s role in phagocytosing abnormal germ cells and residual bodies during spermatogenesis (Oliveira and Alves, 2015). Taken together, our findings provide further support for the key role of Sertoli cells in determining testis mass, sperm production and quality in ungulates (Pintus *et al*., 2015b; Ros-Santaella *et al*., 2019).

Our results show that the investment in secondary sexual characters’ growth is not linked to sperm size but rather to the homogeneity in sperm cell dimension. Findings from comparative studies in this taxonomic group have been so far elusive with both negative (Ferrandiz-Rovira *et al*., 2014) or no (Lüpold *et al*., 2015) relationships between horn/antler development and sperm size. Under this perspective, our findings highlight the relevance of intraspecific studies as they may provide a better understanding of patterns observed across animal species and reveal the signatures of selection (Kleven *et al*., 2008). In red deer, for instance, reduced intramale variation in sperm size is associated with enhanced sperm motility and normal sperm morphology (Ros-Santaella *et al*., 2015), which predict male fertility in this species (Malo *et al*., 2005b). In agreement with a previous study (Ros-Santaella *et al*., 2015), our findings also confirm that normal sperm morphology is associated with reduced intramale variation in sperm length. A smaller within-male CV in sperm size has also been associated with the intensity of sperm competition and testicular investment (Šandera *et al*., 2013; Ros-Santaella *et al*., 2015). The lack of significant relationships between intramale variation in sperm size and spermatogenic function in the common elands raise the possibility that other factors, rather than spermatogenesis *per se*, might play a role in the production of a more uniform sperm population. For instance, in the male great tits (*Parus major*), within-male variation in sperm length is associated with a health-related (haematological) trait but not with male ornaments (Svobodová *et al*., 2018). Another factor that can influence the variation in sperm size is the epididymal environment as sperm morphometry has shown to vary throughout the epididymal regions (Gutiérrez-Reinoso *et al*., 2016), possibly because of fluid reabsorption, hence increased osmolality. In cats, for instance, reduced within-male variation in sperm head ellipticity is related to higher epididymal mass and sperm concentration (Pintus *et al*., 2021). In agreement with these findings, we found that low intramale CV in sperm head ellipticity was associated with higher sperm concentration in the epididymal tails of common elands, a finding that may support the role of the epididymal environment in determining a more uniform sperm population.

Overall, the positive relationship between male secondary sexual characters and spermatogenic function found in this study provides support, at least to some extent, to the fertility-linked hypothesis, which predicts that male phenotype covaries with functional fertility (Sheldon, 1994). In contrast, sperm competition games assume a trade-off between pre- and post-copulatory investment (Parker *et al*., 2013). Across the animal kingdom, evidence has been found to support each or none of these hypotheses (reviewed by Simmons *et al*., 2017). A complex variety of factors may affect the slope of relationships including genetic variation, mating strategy or environmental conditions. In mammals, for instance, the slope of the relationship between primary and secondary sexual characters can be influenced by the mating strategy, as the trade-off is prominent in species in which weaponry investment is effective in female monopolization (Lüpold *et al*., 2014). Common elands usually live in large groups (up to 500 animals) that are often composed of mixed-sex herds of one to four adult males and up to more than 50 females (Bro-Jørgensen *et al*., 2015; Pappas, 2002). Under such mating system in which males likely fail in female monopolization, it would be more advantageous for males to equally invest their limited resources both in pre- and post-copulatory sexual traits. This would contribute to explain the positive relationship between horn size and testicular function found in this study. Another aspect to consider is that in many sexually dimorphic ungulates, secondary sexual characters like horns or antlers can simultaneously serve both as armaments and ornaments. By contrast, in a comparative study in primate species, Lüpold *et al*. (2019) found that only those traits that play a role as sexual ornaments trade-off against testes size in contrast to those regarded as weapons. In the light of our findings, future research effort should be directed towards the understanding of other complex signalling traits of male elands such as pelage ornament (e.g., frontal hairbrush size), dewlap droop, and the peculiar sound presumably produced by the carpal joints, known as knee-clicks (Bro-Jørgensen and Beston, 2015; Bro-Jørgensen and Dabelsteen, 2008). Another aspect to bear in mind is that, under our experimental conditions, optimal nutritional intake was guaranteed to all animals both in terms of quantity and quality, which may mask any potential trade-off between somatic and testicular investment (Parker, 2016). As previously observed in fallow deer, nutrition has a clear impact on spermatogenic function and sperm morphometry during the sexual development (Ros-Santaella *et al*., 2019). It remains therefore to be tested whether under restricted access to nutritional resources a positive relationship would be still found between the development of primary and secondary sexual traits. It is important to highlight that our study was performed on a captive population of common eland. Such experimental design allowed us to explore the relationship between the development of primary and secondary sexual characters under controlled environmental conditions but limits the extrapolation of our findings to natural populations.

The mean horn length found in our study agrees with data reported by previous authors in the adult male common eland (Bro-Jørgensen and Dabelsteen, 2008; Bro-Jørgensen, 2007), which provides support that final horn length is achieved relatively early during the male life and decreases through adulthood because of wearing (Jeffery and Hanks, 1981). Thus, adverse conditions experienced during early development may affect not only the ongoing growth of the individual but also its sexual attractiveness during adulthood (Lindström, 1999). As we could not control for age in our experimental design since this factor is intrinsically linked to sexual development, we cannot rule out that it may have influenced our results. Nevertheless, an unexpected finding of this study was that horn size is not related to the spermatic index nor the percentage of sperm cells with normal morphology, which typically increase throughout the male sexual development (Brito, 2021; Rajak *et al*., 2014). Among the traits employed in this study to assess the horn development, it is remarkable to note that the spiral length showed the strongest correlation with the Sertoli cell efficiency. In addition, we also found that horn basal circumference was significantly associated with intramale variation in sperm head size and testicular investment in the seminiferous tubular and epithelial areas. The spiral length and the basal circumference are the horn traits responsible for the sexual dimorphism in common elands as males show horns with more pronounced spirals and greater basal circumference than females (Pappas, 2002; Jeffery and Hanks, 1981). The spiral ridge of the eland horn core has shown different mechanical and chemical properties compared to other parts of this permanent appendage, which may avoid fractures during male-male contests (Cappelli *et al*., 2018). The spiral of the horn, as bumps and ridges on the horns of various antelopes and gazelles, serve indeed to hold the horns together during the match, allowing the opponents to develop full strength wrestling engagement (Geist, 1966). In a comparative study across 104 bovid species, Caro *et al*. (2003) found that, at least in females, twisted horns are associated with wrestling behaviour. In the same study, the authors also found that the outward-facing and twisted horns of bovids are also associated with polygynous and large mating groups. Such finding was later supported by Bro-Jørgensen (2007) who found that in polygynous mating system, in which sexual selection is likely to be strong, horn shape is more variable and elaborate than in monogamous species, in which horns tend to be simple and straight. Taken together, male elands with more developed spirals and greater basal circumference may signal not only their fighting ability but also their reproductive competence (i.e., spermatogenic efficiency and sperm size homogeneity). Another non-mutually exclusive hypothesis is that increased horn size may provide a better mechanism for thermoregulation, which may also have beneficial effect on spermatogenesis. Within the thermal neutral zone, the common eland metabolism is estimated to be around 30% higher than that of Hereford cattle, a bovid species of comparable size (Taylor and Lyman, 1967), although other authors did not find support to this finding (Kotrba *et al*., 2007). Spermatogenesis requires a cool environment (usually 2-6 °C below the body core temperature), which is normally guaranteed by the scrotal sac and the complex anatomical structures associated with the latter (i.e., pampiniform plexus, dartos, cremaster, and sweat glands). Because these complex anatomical structures develop with age and influence sperm motility and morphology (Brito *et al*., 2012), their role is likely more critical in young individuals because of their higher metabolism compared to that of adults. In line with our hypothesis, in the male Alpine ibex (*Capra ibex*), the early development of horns is a suitable predictor of individual reproductive success later in life, whereas the late development of this secondary sexual trait did not seem to significantly relate to it (Willisch *et al*., 2015). In addition, like other large herbivores adapted to xeric environment (Hetem *et al*., 2016), elands can employ adaptive heterothermy by increasing their body temperature in response to high environmental heat load to reduce the water loss (Taylor and Lyman, 1967; but see also Fuller *et al*., 2004).

Because horn size and shape can provide a more efficient mechanism for heat loss (Picard et al., 1999, 1994), and the latter can be beneficial for the spermatogenesis, we can speculate that bigger horn size can provide a more efficient mechanism not only for the body but, indirectly, also for the testicular thermoregulation. Further studies are required to test whether the development of horn size may contribute to a better testicular thermoregulation. On the other hand, no direct link between horn asymmetry and reproductive function was found, probably because of the absence of environmental stressors in our animal population. Using a large dataset, Chirichella *et al*. (2020) found that environmental stressors (e.g., snow cover duration, population density) experienced in early life influence the symmetrical horn development in the Alpine chamois (*Rupicapra rupicapra*).

In conclusion, greater male investment in secondary sexual characters is related to enhanced spermatogenic function and increased homogeneity in sperm cell dimension. During the sexual development, common elands that display bigger horns show increased testes size, spermatogenic investment, Sertoli cell efficiency and reduced intramale variation in sperm head size and shape. The findings from this study increase our understanding about the role of secondary sexual traits that, beyond sexual selection dynamics, might be involved in the thermoregulatory mechanisms that guarantee the optimal conditions for sperm production.

## Ethics statement

The Czech University of Life Sciences’ eland farm is accredited as research facility according to European and Czech laws for ethical use of animals in research (permits no. 58176/2013-MZE-17214 and no. 63479/2016-MZE-17214). All slaughter process was carried out under the supervision of a state veterinarian according to EU and national legislation and conditions for farm animals (slaughter permit no. SVS/WS22/2012-KVSS and no. SVS/2015/077267-S).

## Authors contributions

EP: Conceptualization, Data curation, Formal analysis, Funding acquisition, Investigation, Methodology, Supervision, Validation, Writing–original draft. RK: Data curation, Resources, Writing–review and editing. JLR-S: Conceptualization, Data curation, Formal analysis, Funding acquisition, Investigation, Methodology, Project administration, Supervision, Validation, Writing–review and editing.

## Funding

This work was supported by CIGA 20145001 (Czech University of Life Sciences, Prague, Czech Republic).

## Conflict of Interest

The authors declare that the research was conducted in the absence of any commercial or financial relationships that could be construed as a potential conflict of interest.

## Permission to reuse and Copyright

Not applicable.

## Notes

### Competing Interest Statement

The authors have declared no competing interest.

## References

Bartoň, L., Bureš, D., Kotrba, R., Sales, J. (2014). Comparison of meat quality between eland (*Taurotragus oryx*) and cattle (*Bos taurus*) raised under similar conditions. Meat Sci. 96, 346–352. doi: 10.1016/j.meatsci.2013.07.016

Benítez, H. A., Lemic, D., Villalobos-Leiva, A., Bažok, R., Órdenes-Claveria, R., Pajač Živković, I., Mikac, K. M. (2020). Breaking symmetry: Fluctuating asymmetry and geometric morphometrics as tools for evaluating developmental instability under diverse agroecosystems. Symmetry 12, 1789. doi: 10.3390/sym12111789

Brito, L. F. C. (2021). Sexual development and puberty in bulls. In Bovine reproduction, John Wiley & Sons, NJ, USA, pp 58–78.

Brito, L. F., Barth, A. D., Wilde, R. E., Kastelic, J. P. (2012). Testicular vascular cone development and its association with scrotal temperature, semen quality, and sperm production in beef bulls. Anim. Reprod. Sci. 134, 135–140. doi: 10.1016/j.anireprosci.2012.08.025.

Bro-Jørgensen, J., and Beeston, J. (2015). Multimodal signalling in an antelope: fluctuating facemasks and knee-clicks reveal the social status of eland bulls. Anim. Behav. 102, 231–239. doi: 10.1016/j.anbehav.2015.01.027.

Bro-Jørgensen, J., and Dabelsteen, T. (2008). Knee-clicks and visual traits indicate fighting ability in eland antelopes: multiple messages and back-up signals. BMC Biology 6, 1–8. doi: 10.1186/1741-7007-6-47.

Bro-Jørgensen, J. (2007) The intensity of sexual selection predicts weapon size in male bovids. Evolution 61,1316–1326. doi: 10.1111/j.1558-5646.2007.00111.x.

Budaev, S. V. (2010). Using principal components and factor analysis in animal behaviour research: Caveats and guidelines. Ethology 116, 472–480. doi: 10.1111/j.1439-0310.2010.01758.x.

Cappelli, J., García, A. J., Kotrba, R., Gambin Pozo, P., Landete-Castillejos, T., Gallego, L., Ceacero, F. (2018). The bony horncore of the common eland (*Taurotragus oryx*): composition and mechanical properties of a spiral fighting structure. J. Anat. 232, 72–79. doi: 10.1111/joa.12708.

Caro, T. M., Graham, C. M., Stoner, C. J., Flores, M. M. (2003). Correlates of horn and antler shape in bovids and cervids. Behav. Ecol. Sociobiol. 55, 32–41. doi: 10.1007/s00265-003-0672-6.

Chirichella, R., Rocca, M., Brugnoli, A., Mustoni, A., Apollonio, M. (2020) Fluctuating asymmetry in Alpine chamois horns: An indicator of environmental stress. Evol. Ecol. 34, 573–587. doi: 10.1007/s10682-020-10051-3.

Dines, J. P., Mesnick, S. L., Ralls, K., May-Collado, L., Agnarsson, I., Dean, M. D. (2015). A trade-off between precopulatory and postcopulatory trait investment in male cetaceans. Evolution. 69, 1560–1572. doi: 10.1111/evo.12676.

Ditchkoff, S. S., Lochmiller, R. L., Masters, R. E., Hoofer, S. R., Bussche, R. A. V. D. Major-histocompatibility-complex-associated variation in secondary sexual traits of white-tailed deer (*Odocoileus virginianus*): evidence for good-genes advertisement. Evolution 55, 616–25 (2001). doi: 10.1554/0014-3820(2001)055[0616:mhcavi]2.0.co;2.

Doyle, J. M. (2011). Sperm depletion and a test of the phenotype-linked fertility hypothesis in gray treefrogs (*Hyla versicolor*). Can. J. Zool. 89, 853–858. doi: 10.1139/Z11-060.

Ezenwa, V. O., Jolles, A. E. (2008). Horns honestly advertise parasite infection in male and female African buffalo. Anim. Behav. 75, 2013–2021. doi: 10.1016/j.anbehav.2007.12.013.

Ferrandiz-Rovira, M., Lemaître, J. F., Lardy, S., López, B.C. Cohas, A. (2014). Do pre-and post-copulatory sexually selected traits covary in large herbivores? BMC Evol. Biol. 14, 79. doi: 10.1186/1471-2148-14-79.

Fuller, A., Maloney, S. K., Mitchell, G., Mitchell, D. (2004). The eland and the oryx revisited: body and brain temperatures of free-living animals. In International congress series 1275, 275– 282).

Geist, V. (1966). The evolution of horn-like organs. Behaviour 27, 175–214.

Glazier, D.S. (2022). Complications with body-size correction in comparative biology: possible solutions and an appeal for new approaches. J Exp Biol. 225, jeb243313. doi: 10.1242/jeb.243313.

Gutiérrez-Reinoso, M. A., García-Herreros, M. (2016). Normozoospermic versus teratozoospermic domestic cats: differential testicular volume, sperm morphometry, and subpopulation structure during epididymal maturation. Asian J. Androl. 18,871–878. doi: 10.4103/1008-682X.187583.

Hetem, R. S., Maloney, S. K., Fuller, A., Mitchell, D. (2016). Heterothermy in large mammals: inevitable or implemented? Biol. Rev. 91, 187–205. doi: 10.1111/brv.12166.

IUCN. The IUCN red list of threatened species (2024). https://www.iucnredlist.org/species/22055/115166135

Jeffery, R. C. V., and Hanks, J. (1981). Age determination of eland Taurotragus oryx (Pallas, 1766) in the Natal Highveld. African Zoology 16, 113–122.

Kekäläinen, J., Pirhonen, J., Taskinen, J. (2014). Do highly ornamented and less parasitized males have high quality sperm? - an experimental test for parasite-induced reproductive trade-offs in European minnow (*Phoxinus phoxinus*). Ecol Evol. 4, 4237–4246. doi: 10.1002/ece3.1267.

Kleven, O., Laskemoen, T., Fossøy F., Robertson, R.J., Lifjeld, J. T. (2008). Intraspecific variation in sperm length is negatively related to sperm competition in passerine birds. Evolution 62, 494–499. doi: 10.1111/j.1558-5646.2007.00287.x.

Kotrba, R., Knížková, I., Kunc, P., Bartoš, L. (2007). Comparison between the coat temperature of the eland and dairy cattle by infrared thermography. J. Therm. Biol. 32, 355–359. doi: 10.1016/j.jtherbio.2007.05.006.

Kruuk, E. B., Slate, J., Pemberton, J. M., Brotherstone, S., Guinness, F., Clutton-Brock T. (2002). Antler size in red deer: heritability and selection but no evolution. Evolution. 56,1683–1695. doi: 10.1111/j.0014-3820.2002.tb01480.x.

Lindström, J. (1999). Early development and fitness in birds and mammals. Trends Ecol. Evol. 14, 343–348. 10.1016/S0169-5347(99)01639-0

Lundrigan, B. (1996). Morphology of horns and fighting behavior in the family Bovidae. J. Mammal. 77, 462–475. doi: 10.2307/1382822.

Lüpold, S., Simmons, L. W., Grueter, C. C. (2019). Sexual ornaments but not weapons trade off against testes size in primates. Proc. R. Soc. Lond. Ser. B-Biol. Sci., 286, 20182542. doi: 10.1098/rspb.2018.2542.

Lüpold, S., Simmons, L. W., Tomkins, J. L., Fitzpatrick, J. L. (2015). No evidence for a trade-off between sperm length and male premating weaponry. J. Evol. Biol. 28, 2187–2195. doi: 10.1111/jeb.12742.

Lüpold, S., Tomkins, J. L., Simmons, L. W., Fitzpatrick, J. L. (2014). Female monopolization mediates the relationship between pre-and postcopulatory sexual traits. Nat. Commun. 5, 1–8. doi: 10.1038/ncomms4184.

Malo, A. F., Roldan, E. R., Garde, J., Soler, A. J., Gomendio, M. (2005a). Antlers honestly advertise sperm production and quality. Proc. Biol. Sci. 272, 149–57. doi: 10.1098/rspb.2004.2933.

Malo, A. F., Garde, J. J., Soler, A. J., García, A. J., Gomendio, M., Roldan, E. R. (2005b). Male fertility in natural populations of red deer is determined by sperm velocity and the proportion of normal spermatozoa. Biol. Reprod. 72,822–829. doi: 10.1095/biolreprod.104.036368.

Mautz, B. S., Møller, A.P., Jennions, M. D. (2013). Do male secondary sexual characters signal ejaculate quality? A meta-analysis. Biol. Rev. Camb. Philos. Soc. 88, 669–682. doi: 10.1111/brv.12022.

Navara, K. J., Anderson, E. M., Edwards, M. L. (2012). Comb size and color relate to sperm quality: a test of the phenotype-linked fertility hypothesis. Behav. Ecol. 23, 1036–1041. doi: 10.1093/beheco/ars068.

O’Donnell, L., Smith, L. B., Rebourcet, D. (2022). Sertoli cells as key drivers of testis function. Semin. Cell Dev. Biol. 121, 2–9. doi: 10.1016/j.semcdb.2021.06.016.

Oliveira, P.F. and Alves, M.G. (2015). The Sertoli cell at a glance. In Sertoli cell metabolism and spermatogenesis (Springer Publisher).

Palmer, A. R. (1994). Fluctuating asymmetry analyses: A primer. In Developmental instability: Its origins and evolutionary implications. (Kluwer).

Pappas, L. A. (2002). Taurotragus oryx. Mammalian species 689, 1–5.

Parker, G. A. (2016). The evolution of expenditure on testes. J. Zool. 298, 3–19. doi: 10.1111/jzo.12297.

Parker, G. A., Lessells, C. M., Simmons, L. W. (2013). Sperm competition games: a general model for precopulatory male–male competition. Evolution 67, 95–109. 10.1111/j.1558-5646.2012.01741.x.

Paschoal, L. R. P. and Zara, F. J. (2022). Is there a trade-off between sperm production and sexual weaponry in the Amazon River prawn Macrobrachium amazonicum (Heller, 1862)? Zoology 153, 126029. doi: 10.1016/j.zool.2022.126029.

Pennington, P. M. (2009). Characterization of the common eland (*Taurotragus oryx*) estrous cycle. LSU Master’s Theses 1542. doi: 10.1071/RDv21n1Ab164.

Picard, K., Thomas, D. W., Festa-Bianchet, M., Belleville, F., Laneville, A. (1999). Differences in the thermal conductance of tropical and temperate bovid horns. Ecoscience 6, 148–158. doi: 10.1080/11956860.1999.11682515.

Picard, K., Thomas, D. W., Festa-Bianchet, M., Lanthier, C. (1994). Bovid horns: an important site for heat loss during winter? J. Mammal. 75, 710–713. doi: 10.2307/1382520.

Pintus, E., Kadlec, M., Karlasová, B., Popelka, M., Ros-Santaella, J. L. (2021). Spermatogenic activity and sperm traits in post-pubertal and adult tomcats (*Felis catus*): Implication of intra-male variation in sperm size. Cells 10,624. doi: 10.3390/cells10030624.

Pintus, E., Ros-Santaella, J. L., Garde J. J. (2015a). Variation of spermatogenic and Sertoli cell number detected by fine needle aspiration cytology (FNAC) in Iberian red deer during and out of the breeding season. Reprod. Fertil. Dev. 27, 812–822. doi: 10.1071/RD13419.

Pintus, E., Ros-Santaella, J.L., Garde, J. J. (2015b). Beyond testis size: Links between spermatogenesis and sperm traits in a seasonal breeding mammal. PLoS One 10, e0139240. doi: 10.1371/journal.pone.0139240.

Pintus, E., Ros-Santaella, J.L., Garde J. J. (2014). Diagnostic value of fine needle aspiration cytology in testicular disorders of red deer (*Cervus elaphus*): a case report. J. Wildl. Dis. 50,994– 997. doi: 10.7589/2013-11-304.

Preston, B. T., Stevenson, I. R., Pemberton, J. M., Coltman, D. W., Wilson K. (2003). Overt and covert competition in a promiscuous mammal: the importance of weaponry and testes size to male reproductive success. Proc. Biol. Sci. 270, 633–640. doi: 10.1098/rspb.2002.2268.

Rajak, S. K., Kumaresan, A., Gaurav, M. K., Layek, S. S., Mohanty, T. K., Muhammad Aslam, M. K., Tripathi, U. K., Prasad, S., De, S. (2014). Testicular cell indices and peripheral blood testosterone concentrations in relation to age and semen quality in crossbred (holstein friesian×tharparkar) bulls. Asian-Australas. J. Anim. Sci. 27, 1554–61. 10.5713/ajas.2014.14139.

Ramm, S.A. and Schärer, L. (2014). The evolutionary ecology of testicular function: size isn’t everything. Biol. Rev. Camb. Philos. Soc. 89,874–888. doi: 10.1111/brv.12084.

Rogell, B., Dowling, D.K., Husby, A. (2019). Controlling for body size leads to inferential biases in the biological sciences. Evol Lett. 4, 73–82. doi: 10.1002/evl3.151.

Rogers, D. W., Denniff, M., Chapman, T., Fowler, K., Pomiankowski. A. (2008). Male sexual ornament size is positively associated with reproductive morphology and enhanced fertility in the stalk-eyed fly *Teleopsis dalmanni*. BMC Evol. Biol. 8, 236. doi: 10.1186/1471-2148-8-236.

Roldan, E. R., Cassinello, J., Abaigar, T., Gomendio, M. (1998). Inbreeding, fluctuating asymmetry, and ejaculate quality in an endangered ungulate. Proc. Biol. Sci. 265, 243–248. doi: 10.1098/rspb.1998.0288.

Ros-Santaella, J. L., Kotrba, R., Pintus, E. (2019). High-energy diet enhances spermatogenic function and increases sperm midpiece length in fallow deer (*Dama dama*) yearlings. R. Soc. Open Sci. 6,181972. doi: 10.1098/rsos.181972.

Ros-Santaella, J. L., Pintus, E., Garde, J. J. (2015). Intramale variation in sperm size: functional significance in a polygynous mammal. PeerJ 3, e1478. doi: 10.7717/peerj.1478.

Ros-Santaella, J. L., Domínguez-Rebolledo, A. E., Garde, J. J. (2014). Sperm flagellum volume determines freezability in red deer spermatozoa. PLoS One 9, e112382. doi: 10.1371/journal.pone.0112382.

Šandera, M., Albrecht, T., Stopka P. (2013). Variation in apical hook length reflects the intensity of sperm competition in murine rodents. PLoS One. 8, e68427. doi: 10.1371/journal.pone.0068427.

Santiago-Moreno, J., Toledano-Díaz, A., Pulido-Pastor, A., Gómez-Brunet, A., López-Sebastián, A. (2007). Horn quality and postmortem sperm parameters in Spanish ibex (*Capra pyrenaica hispanica*). Anim. Reprod. Sci. 99, 54–62. doi: 10.1016/j.anireprosci.2006.06.004.

Sheldon, B. C. (1994). Male phenotype, fertility, and the pursuit of extra-pair copulations by female birds. Proc. R. Soc. Lond. Ser. B-Biol. Sci. 257, 25–30. doi: 10.1098/rspb.1994.0089.

Simmons, L. W., Lüpold, S., Fitzpatrick, J. L. (2017). Evolutionary trade-off between secondary sexual traits and ejaculates. Trends Ecol. Evol. 32, 964–976. doi: 10.1016/j.tree.2017.09.011.

Snook, R. R., Gidaszewski, N. A., Chapman, T., Simmons, L. W. (2013). Sexual selection and the evolution of secondary sexual traits: sex comb evolution in Drosophila. J. Evol. Biol. 26, 912–918. doi: 10.1111/jeb.12105.

Streiner, D. L. (2015). Best (but oft-forgotten) practices: the multiple problems of multiplicity-whether and how to correct for many statistical tests. Am. J. Clin. Nutr. 102, 721–728. 10.3945/ajcn.115.113548.

Svobodová, J., Bauerová, P., Eliáš, J., Velová, H., Vinkler, M., Albrecht, T. (2018). Sperm variation in great tit males (*Parus major*) is linked to a haematological health-related trait, but not ornamentation. J. Ornithol. 159, 815–822. doi: 10.1007/s10336-018-1559-7.

Taylor, C. R. and Lyman, C. P. (1967). A comparative study of the environmental physiology of an East African antelope, the eland, and the Hereford steer. Physiol. Zool. 40, 280–295.

Vágner, J. A. (1974). The capture and transport of African animals. International Zoo Yearbook 14, 69–73.

Varea-Sánchez, M., Gómez Montoto, L., Tourmente, M., Roldan, E.R. (2014). Postcopulatory sexual selection results in spermatozoa with more uniform head and flagellum sizes in rodents. PLoS One 9, e108148. doi: 10.1371/journal.pone.0108148.

Vea, I. M. and Shingleton, A.W. (2021). Network-regulated organ allometry: The developmental regulation of morphological scaling. Wiley Interdiscip Rev Dev Biol. 10, e391. 10.1002/wdev.391

Willisch, C. S., Biebach, I., Marreros, N., Ryser-Degiorgis, M. P., Neuhaus, P. (2015). Horn growth and reproduction in a long-lived male mammal: no compensation for poor early-life horn growth. Evol. Biol. 42, 1–11. doi: 10.1007/s11692-014-9294-3.

